# rCASC: reproducible Classification Analysis of Single Cell sequencing data

**DOI:** 10.1101/430967

**Authors:** Luca Alessandrì, Marco Beccuti, Maddalena Arigoni, Martina Olivero, Greta Romano, Gennaro De Libero, Luigia Pace, Francesca Cordero, Raffaele A Calogero

## Abstract

**Summary:** Single-cell RNA sequencing has emerged as an essential tool to investigate cellular heterogeneity, and highlighting cell sub-population specific signatures. Nowadays, dedicated and user-friendly bioinformatics workflows are required to exploit the deconvolution of single-cells transcriptome. Furthermore, there is a growing need of bioinformatics workflows granting both functional, i.e. saving information about data and analysis parameters, and computation reproducibility, i.e. storing the real image of the computation environment. Here, we present rCASC a modular RNAseq analysis workflow allowing data analysis from counts generation to cell sub-population signatures identification, granting both functional and computation reproducibility.

**Availability and Implementation:** rCASC is part of the reproducible bioinfomatics project. rCASC is a docker based application controlled by a R package available at https://github.com/kendomaniac/rCASC.

**Supplementary information:** Supplementary data are available at *rCASC github*

## 1 Introduction

Single cell analysis is instrumental to understand the functional differences existing between cells within a tissue. Individual cells of the same phenotype are commonly viewed as identical functional units of a tissue or organ. However, published single cells sequencing results (Buettner, et al., 2015) suggest the presence of a complex organization of heterogeneous cell states producing together system-level functionalities. Single cell analysis focuses on the understanding differences characterizing any cell within a population of cells. A mandatory element of single cell RNAseq is the availability of dedicated bioinformatics workflows. In the last few years a lot of tools have been developed for the identification of tissue cell subpopulations (Hwang, et al., 2018). However, sub-population identification might require some preprocessing steps of the single-cell sequencing data, depending on the technology in use and on some specific cell characteristics, e.g. cell state. Furthermore, after cell partitioning, extra steps are required to identify cell sub-population signatures, e.g. markers identification. rCASC is a modular workflow, based on docker technology, allowing processing of 10XGenomics, inDrop and whole transcripts single-cell sequences from fastq to the definition of cell sub-population signatures. Furthermore, rCASC addresses the problem of functional and computational reproducibility, which is becoming a very important topic, because of the “Data Reproducibility Crisis” (Allison, et al., 2018).

**Figure 1:**
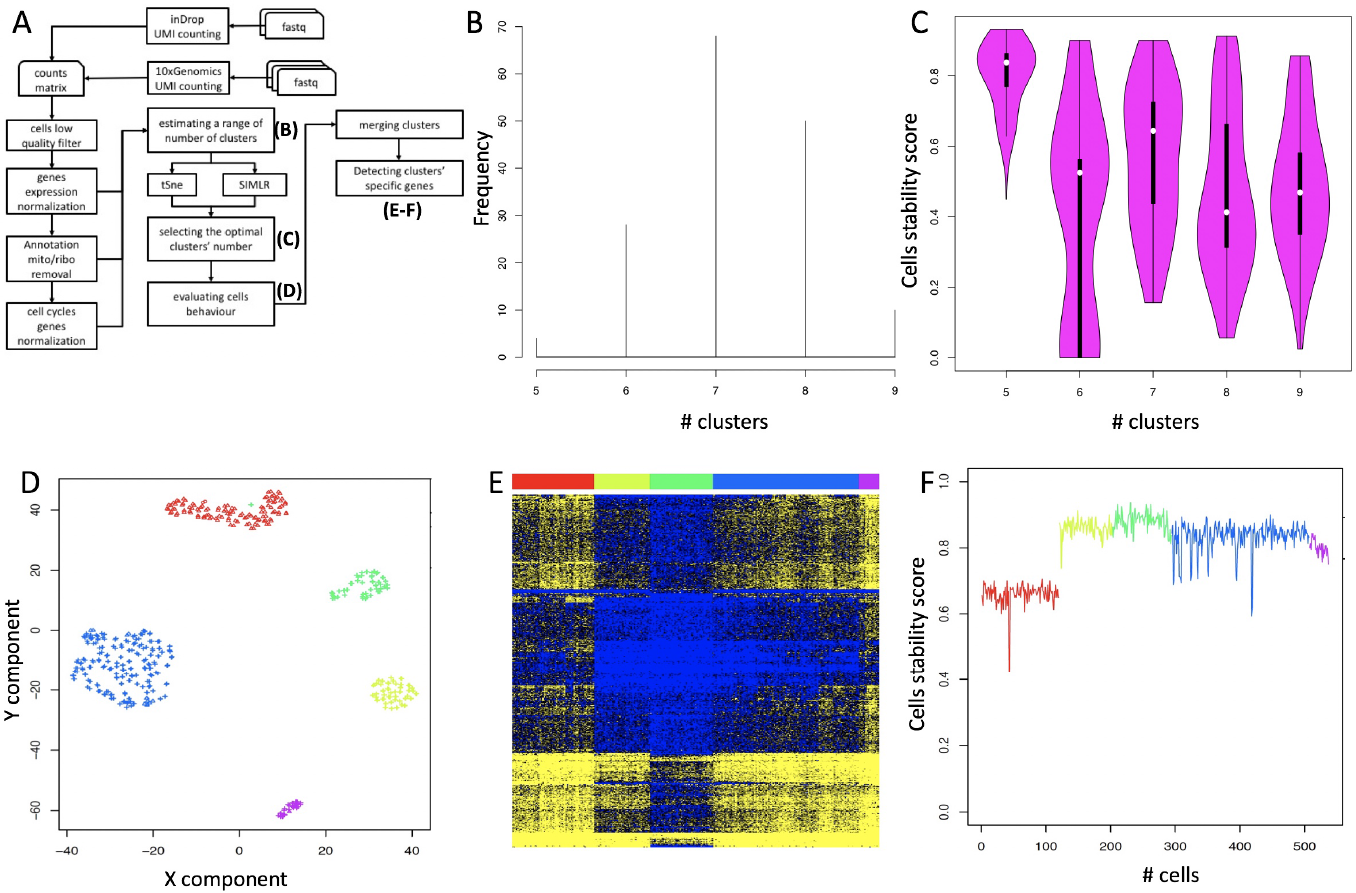
rCASC workflow. A) rCASC modules, outputs for the relevant steps of the workflow are shown in capital letters in parenthesis. B) Depicting the range of clusters to be investigated with k-mean clustering tools (SIMLR or tSne). C) Cell stability score for the selected range of clusters. D) SIMLR clusterization results for the number of clusters showing the most homogeneous cell stability score, i.e. 5 clusters. E) Z-score heatmap of prioritized clusters-specific genes. Color bar refers to clusters colors in D. F) Cell stability score in E. Colors refers to clusters’ colors.

## 2 Methods

rCASC is developed within the umbrella of the Reproducible Bioinformatics Project (*www.reproducible-bioinformatics.org*), which is an open-source community aiming to develop reproducible bioinformatics workflows. Each module of rCASC is implemented in a docker container, and it is compliant with the rules proposed by Sandve (Sandve, et al., 2013) to guarantee reproducibility. The key elements of rCASC workflow are shown in Fig. 1A, and the main functionalities are summarized below.

### Data preprocessing

rCASC allows processing of fastq derived by 10XGenomics and inDrop platforms to generate a cell count matrix annotated using ENSEMBL gene model (Supplementary Section 2). Furthermore, counts matrix, using ENSEMBL gene model, can be processed within rCASC. The most relevant preprocessing modules of rCASC (Supplementary Section 3) allow visualization of the numbers of genes detected in each cell with respect to the cells total reads, removal of low quality cells using Lorenz statistic (Diaz, et al., 2016), removal of ribosomal and mitochondrial genes and the association of gene symbol to the ENSEMBL gene identifier, data normalization (Bacher, et al., 2017), detection of possible cell cycle bias (Liu, et al., 2017) and removal of such effects from the data (Barron and Li, 2016).

### Cell heterogeneity analysis

The optimal number of cells partitions is detected inducing perturbations in the structure of the cell data set, i.e. removing a random subset of cells and repeating the clustering. The rational of this approach is that a robust cluster of cells should contain the same set of cells independently by the perturbation of the overall dataset. The bootstrapped dataset is analyzed with a graph-based community detection method (https://github.com/ppapasaikas/griph), allowing the identification of the range of number of clusters observable perturbing the cells dataset structure (Fig. 1B, Supplementary Section 4). Then, the range of number of clusters is probed using SIMLR (Wang, et al., 2017), a clustering framework learning a sample-to-sample similarity measure from expression data. A cell stability score (Supplementary Section 5), indicating the fraction of bootstraps in which a cell is allocated in a specific cluster, is used to identify the optimal number of clusters for the cell sub-populations representation (Fig. 1C). Cells are then plotted in each cluster with a specific symbol indicating its stability (Fig. 1D). Furthermore, the shuffling of unstable cells between nearby clusters can be visualized in a video in which each bootstrap is a frame of a video.

### Clusters specific feature selection

The identification of clusters specific signatures is addressed with two different methods (Supplementary Section 6). The ANOVA-like method from edgeR (Robinson, et al., 2010) is used in case of the presence a reference cluster, e.g. in a cells activation experiment it could be the cluster of resting cells undergoing to activation/differentiation by an external stimulus. In case a reference cluster is not available SIMLR (Wang, et al., 2017) provides a gene prioritization, measuring how gene expression values across cells correlate with the learned cell-to-cell similarity. This information combined with dataset bootstraps allows the identification of genes which are the main players in clusters organization. The genes selected with the above-mentioned approaches can be then visualized with a supervised heatmap ordering cells according to the belonging cluster (Fig. 1E). The cell stability in each cluster is also provided (Fig. 1F). GUI: Implementation of rCASC functions within *4SeqGUI* is in progress, to make the analysis workflow user-friendly and suitable for users lacking of scripting knowledge.

## Results

The main objective of rCASC is the identification of the most robust partitioning of cell sub-populations within a reproducible framework. The comparison of rCASC with four single-cell analysis workflows (Supplementary Section 8) indicate that rCASC provides unique features, e.g. jackknife resampling for cluster robustness evaluation. The cluster’s robustness, evaluated measuring the persistence of cells in a cluster, as consequence of jackknife resampling, provides a better estimation of clusters stability with respect to other measurements as the silhouette plot (Supplementary Section 5, Fig. 23 A,B). With respect to other workflows, rCASC uses as clustering tool SIMLR, which was shown to outer-performed at least part of the methods implemented in other workflows. rCASC modularity structure easily allows the implementation of other pre/post processing methods and supports the implementation of other clustering methods within the resampling framework. Furthermore, rCASC is the only workflow granting functional and computational reproducibility.

## Conclusion

In conclusion, rCASC is a workflow with valuable new features that could help researchers in defining cells sub-populations and detecting sub-population specific markers, under the umbrella of data reproducibility.

## References

Allison, D.B., Shiffrin, R.M. and Stodden, V. Reproducibility of research: Issues and proposed remedies. Proceedings of the National Academy of Sciences of the United States of America 2018;115(11):2561–2562.

Bacher, R., et al. SCnorm: robust normalization of single-cell RNA-seq data. Nature methods 2017;14(6):584–586.

Barron, M. and Li, J. Identifying and removing the cell-cycle effect from single-cell RNA-Sequencing data. Sci Rep 2016;6:33892.

Buettner, F., et al. Computational analysis of cell-to-cell heterogeneity in single-cell RNA-sequencing data reveals hidden subpopulations of cells. Nature biotechnology 2015;33(2):155–160.

Diaz, A., et al. SCell: integrated analysis of single-cell RNA-seq data. Bioinformatics 2016;32(14):2219–2220.

Hwang, B., Lee, J.H. and Bang, D. Single-cell RNA sequencing technologies and bioinformatics pipelines. Exp Mol Med 2018;50(8):96.

Liu, Z.H., et al. Reconstructing cell cycle pseudo time-series via single-cell transcriptome data. Nature Communications 2017;8.

Robinson, M.D., McCarthy, D.J. and Smyth, G.K. edgeR: a Bioconductor package for differential expression analysis of digital gene expression data. Bioinformatics 2010;26(1):139–140.

Sandve, G.K., et al. Ten simple rules for reproducible computational research. PLoS computational biology 2013;9(10):e1003285.

Wang, B., et al. Visualization and analysis of single-cell RNA-seq data by kernel-based similarity learning. Nature methods 2017;14(4):414–416.

